# Automated Tumor and FUS Lesion Quantification on Multi-frequency Harmonic Motion and B-mode Imaging Using a Multi-modality Neural Network

**DOI:** 10.1101/2024.10.02.616303

**Authors:** Shiqi Hu, Yangpei Liu, Xiaoyue Li, Elisa E. Konofagou

## Abstract

Harmonic Motion Imaging (HMI) is an ultrasound elasticity imaging method that measures the mechanical properties of tissue using amplitude-modulated acoustic radiation force (AM-ARF). By estimating tissue’s on-axis oscillatory motion, HMI-derived displacement images represent localized relative stiffness and can predict the tumor response to neoadjuvant chemotherapy (NACT) and monitor focused ultrasound (FUS) ablation therapy. Multi-frequency HMI (MF-HMI) excites tissue at various AM frequencies simultaneously, which allows for image optimization without prior knowledge of inclusion size and stiffness. However, challenges remain in size estimation as inconsistent boundary effects result in different perceived sizes across AM frequencies. Herein, we developed an automated tumor and FUS lesion quantification method using a transformer-based multi-modality neural network, HMINet. It was trained on 380 pairs of MF-HMI and B-mode images of phantoms and *in vivo* orthotopic breast cancer mice (4T1). Test datasets included phantoms (n = 32), *in vivo* 4T1 mice (n = 24), breast cancer patients (n = 16), and a FUS-induced lesion, with average segmentation accuracy (Dice Similarity Score) of 0.95, 0.86, 0.82, and 0.87, respectively. To increase the generalizability of HMINet, we applied a transfer learning strategy, i.e., fine-tuning the model using patient data. For NACT patients, the displacement ratios (DR) between the tumor and surrounding tissue were calculated based on HMINet-segmented boundaries to predict tumor response based on stiffness changes.

## I. Introduction

Accurate tumor volume assessment is critical for the diagnosis and therapy of breast cancer. Whereas benign lesions are usually associated with distinct, well-circumscribed margins, irregular-shaped masses with spiculate edges are highly suspected of malignancy. These imaging features not only help physicians to identify suspicious masses for biopsy but also infer therapeutic outcomes. In neoadjuvant chemotherapy (NACT), a systemic treatment before surgery to downstage early-stage, locally advanced breast cancer, its response gives guidance to adjuvant treatment and informs prognosis [1]. Image-derived biomarkers can be established to predict patients’ treatment outcomes after NACT, which is evaluated by pathologic complete response (pCR: the absence of cancer cells in the primary tumor and lymph nodes and is associated with long-term clinical benefits such as disease-free survival (DFS) and overall survival (OS) [2]). Early prediction of NACT response is imperative for timely treatment adjustment in clinical practice. In non-invasive thermal ablation therapy using focused ultrasound (FUS), thermal heating causes protein denaturation and lesion formation, and the intraoperative imaging of the affected region allows surgeons to track procedure progress and determine endpoints [3]. A high spatial and temporal-resolution imaging technique guarantees a reliable treatment and facilitates the clinical translation of FUS.

Among all imaging techniques, ultrasound is routinely used for breast imaging because it provides quality images of dense breasts while being non-ionizing, non-invasive, and cost-effective [4]. On traditional B-mode images, masses usually present as darker regions, i.e., hypoechoic, compared with healthy tissues. On the other hand, ultrasound elastography estimates the underlying mechanical properties of tissue to provide insights into its physiological conditions. Tumors are often substantially stiffer than normal tissues due to the desmoplastic reaction, which is featured by the deposition and alteration of extracellular matrix (ECM) proteins [5]. The desmoplasia-associated increased stiffness is considered a critical factor in tumorigenesis, proliferation, and drug resistance in breast cancer [6], [7] and can be captured by elasticity imaging. Though B-mode is well-established for tumor size measurement, its performance can be unsatisfactory with fuzzy tumor edges and mixed echogenicity patterns [8]. The addition of elasticity images can increase the segmentation accuracy [9].

Harmonic Motion Imaging (HMI) is an ultrasound elastography technique. It deforms the tissue using focused acoustic radiation force (ARF) and estimates the resulting on-axis displacements. The term “Harmonic motion” refers to the oscillatory tissue motion induced by amplitude-modulated acoustic radiation force (AM-ARF). Because the pre-defined modulation frequency is distinct from physiologic motion (e.g., breathing) in the low-frequency range, noise that compromises displacement estimation can be efficiently suppressed using bandpass filtering [10]. Compared with shear wave elastography (SWE), which measures off-axis displacements to derive shear modulus quantitatively, and Acoustic Radiation Force Impulse (ARFI) imaging, which measures on-axis displacements after the impulse excitation, HMI uses focused AM-ARF (AM frequency: 25 to 1000 Hz) and measures on-axis displacements during tissue excitation. These allow HMI to be less prone to shear wave reflections and able to penetrate deeper.

So far, our group has demonstrated HMI’s ability in tissue characterization [11], [12], tumor treatment response prediction [13] and FUS treatment monitoring [10], [14]. Whereas MRI can accurately map temperature changes during FUS ablation, it suffers from poor temporal resolution and less accessibility. Alternatively, real-time ultrasound-based monitoring is easier to integrate with FUS and can easily validate that the treatment is targeted at the right window [15]. However, chaotic cavitation events during FUS complicate the estimation of lesion sizes on B-mode images. By generating a high-intensity focused AM-ARF, HMI-guided focused ultrasound (HMIgFUS) ablates the targeted tissue and, in the meantime, assesses the formation of lesions through stiffness-based monitoring. The capability of ablating and monitoring simultaneously is a unique advantage of the HMIgFUS system.

Recently, the development of multi-frequency HMI (MF-HMI) allows the generation of HMI displacements at various AM frequencies simultaneously [16], [17]. It offers more flexibility and can optimize image quality at different frequencies regarding the inclusion dimension and stiffness, as inclusions respond differently to acoustic radiation oscillations. A higher frequency is more sensitive to stiffer and/or smaller inclusions. For stiffer inclusion, the shear wave speed is higher, which is associated with a longer wavelength and a more significant boundary estimation error. Increased AM frequency results in a shorter wavelength and therefore reduces the shearing effects; however, image quality cannot be improved monotonically as the signal-to-noise ratio will be negatively influenced as attenuation also increases with higher shear wave frequencies [17]. This leads to the need to explore MF-HMI images to enhance lesion detectability without prior knowledge of its dimensions and stiffness. Manual segmentation on MF-HMI images can be time-consuming and is influenced by factors such as the operator’s experience and the choice of dynamic range. For the downstream tumor assessment based on HMI images, predictions to NACT are highly subjected to regions of interest (ROI) selection. As a result, an automated segmentation method for MF-HMI can increase the objectivity and accuracy in NACT response prediction and FUS therapy monitoring. A UNet-based segmentation network on MF-HMI was developed in our previous work in which we showed segmentation results on phantom and in vivo murine tumors [18]. However, it’s performance in clinical cases can be further improved.

Many efforts have been made in the machine learning field to facilitate the analysis of multi-parametric ultrasound images for tissue characterization and NACT prediction. Xie et al. [19] employed automated breast ultrasound (ABUS) and contrast-enhanced ultrasound (CEUS) to generate tumor size, blood flow, and vessel information to classify pCR/non-pCR after NACT. Misra et al. [20] proposed a multimodal fusion network for segmentation and malignancy classification utilizing B-mode and strain elastography. In fact, the prediction method based on multi-frequency HMI is worth more exploration, as the stiffness information is intrinsically linked with the tumor microenvironment [21], and MF-HMI provides superior quality compared with other elasticity imaging. For FUS lesion monitoring, Han et al. [22] established an accurate quantitative model for lesion monitoring, which relied on selecting Nagakami parameters and window size. Wu et al [23] proposed a lesion segmentation network for multi-wavelength photoacoustic (MWPA) imaging; but MWPA acquisition can only be implemented before and after FUS treatment. HMI has real-time potential thanks to its hardware configuration, but a precise yet efficient segmentation algorithm is still needed to provide high temporal resolution for FUS monitoring.

In this study, we propose a multi-modality neural network, HMINet, for automated tumor and FUS lesion quantification. Major contributions were made in the following aspects:

1) Constructed a multi-modality neural network, HMINet, with enhanced segmentation accuracy. We further improved its clinical generalizability using transfer learning.
2) Demonstrated that the multimodal strategy is crucial for increasing the segmentation robustness on clinical data, especially for tumors with mixed echogenic patterns.
3) Automated the workflow to predict tumor response to NACT, enabling automatic generation of regions of interest (ROI), which ensures the objectivity of tumor response evaluation.
4) Employed HMINet to segment lesions at different frames acquired by HMIgFUS system during FUS ablation.

To our knowledge, this is the first application of a deep learning-based segmentation method in ROI selection for NACT response prediction and FUS monitoring using elastography.

## II. Methods

### A. Dataset

This study was approved by the Institutional Animal Care and Use Committee and the Institutional Review Board of Columbia University. A total of 380 and 72 pairs of MF-HMI and B-mode images were used for training and validation, including breast-tissue mimicking phantoms of elastic inclusions (N = 214, diameter: 1.7, 2.5, 4.1, 6.5, 10.4 mm, Young’s modulus: 6, 9, 36, 72 kPa, in an 18-kPa background), inclusions (N = 94, diameter: 3, 5, 8, 10, 13 mm, Young’s modulus: 31 - 160 kPa, background: 5-10 kPa) with/without phase aberrations generated by *ex vivo* animal tissue (2-3 cm layers of chicken breast or porcine tissue) or metallic wire mesh, as well as *in vivo* 4T1 murine tumors (N =72, at three different timepoints). 4T1 tumor cell lines (1e^6^), which are used to model human triple-negative breast cancer (TNBC), were subcutaneously implanted into the left lower quadrant abdomen of eight-week-old female BALB/c mice (obtained from The Jackson Laboratory). Tumor progression was monitored weekly using MF-HMI for three weeks, starting from 6 days after tumor cell injection.

The test dataset included a FUS lesion and *in vivo* human malignant tumors (sixteen samples from nine patients with multiple time points) (summarized in Table I). Written consent was obtained before the baseline scan. FUS-induced lesion on freshly obtained *ex vivo* chicken breast tissue was monitored by MF-HMI during ablation.

### B. Multi-frequency Harmonic Motion Imaging

In a conventional HMI setup, a focused ultrasound transducer generates oscillatory ARF for tissue deformation, and a co-aligned imaging transducer estimates the resulting displacements [11], [12]. Oscillatory ARF is achieved by amplitude modulation (AM) using an AM signal containing one or several oscillation frequencies (ranging from 25 to 1000 Hz). In multi-frequency HMI (MF-HMI), multi-frequency excitation (MFE) is achieved by either summing up a set of sinusoids [17] (1) or generating a chirp sequence that contains a range of frequencies (2).

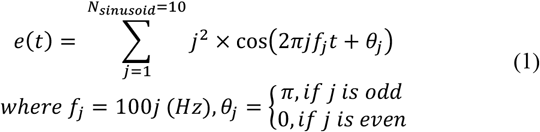

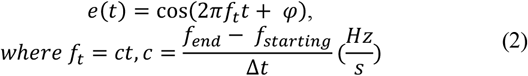

Hossain *et al*. [16] developed the single-transducer HMI (ST-HMI) system that employs an imaging transducer to both excite tissue motion and estimate the resulting displacement. The sequence of MF-ST-HMI is a sum of ten sinusoids (from 100 Hz to 1000 Hz, 100 Hz at a step), as shown in (1). The AM signal was then used to modulate the amplitude (pulse duration) of the excitation pulses. A squared scaling factor, *j* ^2^, was applied to balance more significant acoustic attenuation at high AM frequencies. Multiple short tracking pulses were sent in between long ARF excitation pulses for motion tracking. The overall interleaved excitation-tracking sequence had a frame rate of 10 kHz and a length of 40-60 ms, consisting of 4-6 cycles of the fundamental AM frequency (100 Hz).

The experimental setup for ST-HMI is illustrated in Fig. 1a. The imaging transducer (L7-4 (Philips Healthcare) for phantom and patient studies, L11-5 (Verasonics) for mouse studies) was driven by a Vantage Research System (Vantage 256, Verasonics Inc., Kirkland, WA, USA). Focused excitation and tracking beams were generated by sub-aperture with center frequencies (*f*_*c*_) and F numbers of 4 MHz, 2.25, and 6 MHz, 1.75, respectively. To acquire 2D MF-HMI images, electronic steering was used to translate the sub-aperture from one side to the other, covering a lateral field of view of 18 mm with 32/38 evenly spaced RF lines (0.6 mm spacing). A frame of B-mode with 128 RF lines was collected right after HMI measurements. For clinical data acquisition, a commercially available handheld ultrasound probe (Butterfly) was first used to locate the tumor, following which MF-HMI and research B-mode were acquired after localization using the L7-4 transducer.

**Fig. 1.**
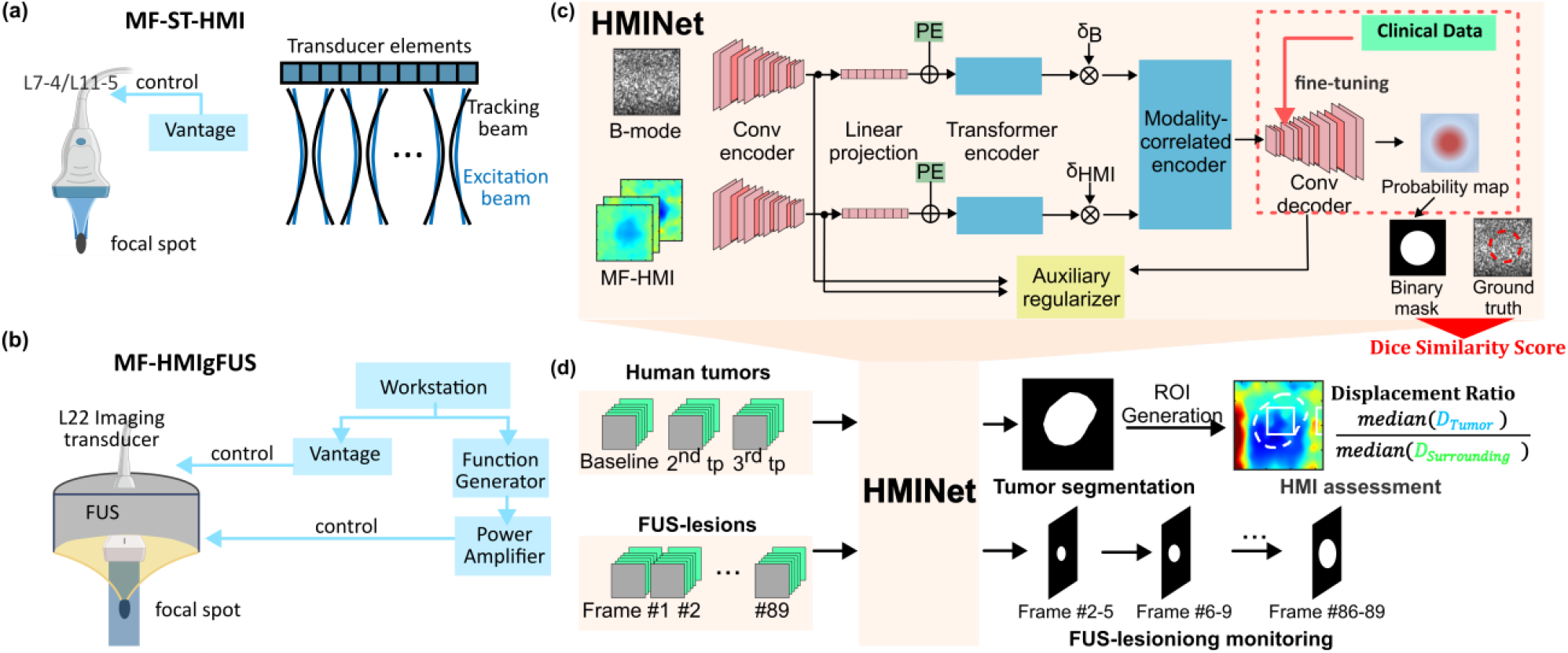
(a): experimental setup of ST-HMI system. (b) experimental setup of HMIgFUS system. (c) HMINet consists of two 5-stage convolutional encoders, two Transformer encoders, a modality-correlated Transformer encoder, and a 5-stage convolutional decoder with channel attention. PE: position encoding for spatial correlation learning. *δ*_*B*_and *δ*_*HMI*_ are Bernoulli indicators designed to increase robustness in the case of missing modality. Transfer learning: *Conv decoder* was re-trained on clinical data. (d) Pipeline for tumor and FUS lesion segmentation and NACT response prediction.

**Fig. 2.**
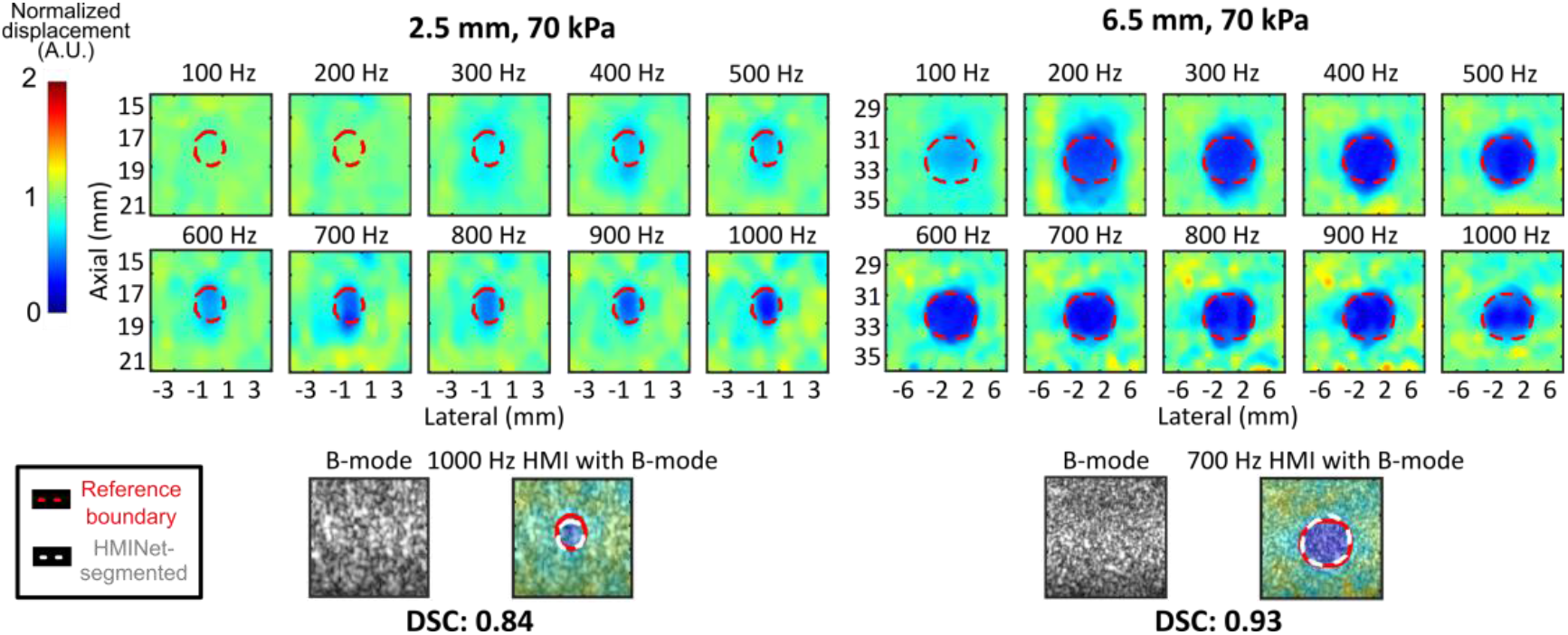
MF-HMI and B-mode images of cylindrical inclusions overlaid with HMINet-segmented boundaries (white) and reference boundaries (red). Images of 2.5 mm, 70 kPa, and 6.5 mm, 70 kPa inclusions are shown.

Fig. 1b shows the experimental setup for HMIgFUS system. It consisted of a single-element focused ultrasound (FUS) transducer (*f*_*c*_ = 4 MHz) driven by a dual-channel arbitrary waveform generator amplified by a 50-dB gain power amplifier and an imaging transducer L22-14vxLF (*f*_*c*_ = 15.625 MHz, Vermon, Tours, France) [24]. It was driven by the Vantage system and was confocally aligned with FUS to track the resulting tissue response. FUS generated prolonged AM-FUS excitation pulses (compared to HMI) for thermal ablation and tracking tissue motion at the same time. The multi-frequency AM signal was a chirp signal with linearly varying instantaneous frequency. The ablation lasted for 120 s with a 50% duty cycle. The imaging frame rate was 10 kHz. FUS ran continuously during the entire 120 s-ablation, while the imaging pulses stopped every 0.15s for data transfer and resumed after the transfer was finished, resulting in a total of 89 MF-HMI acquisitions.

### C. Data Processing

Channel data was beamformed using a GPU-based delay-and-sum (DAS) algorithm. For B-mode image formation, envelope detection and log compression were applied. For HMI data processing, spline interpolation was used on RF data to fill the gaps between RF lines in ST-HMI and recover RF lines affected by FUS interference. Normalized 1D cross-correlation (window size: 2λ, 99% overlap) [25] was applied to estimate either the peak-to-peak displacement or peak positive displacement in the axial direction. Ten displacement images, from 100 to 1000 Hz (100 Hz at a step) or from 150 to 600 Hz (50 Hz at a step), were derived from every ST-HMI or HMIgFUS acquisition. Median filtering (window size of 1.5 mm) was applied to filter out outliers from displacement estimation errors and noise. An axial displacement profile (lateral size: 1.5 mm) was obtained in the background to correct the acoustic force attenuation across depth for ST-HMI images. More details can be found in [17], [24]

Due to the inherent heterogeneity of breast tumors, the relative stiffness of the tumor can vary broadly across patients depending on their pathological conditions. Therefore, we calculated the displacement of the tumor relative to the surrounding tissue to assess the tumor response over the course of NACT treatment. As illustrated in (3), displacement ratios (DR) were computed as the ratio between the median displacement (D) within the tumor region of interest (ROI) and D in the surrounding tissue ROI.

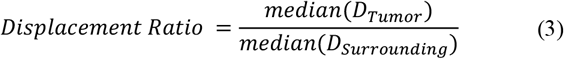

### D. Multi-modality Segmentation Network

In recent years, some pre-trained models that require few or no domain adaptation have shown great promise in medical image segmentation, e.g., segment-anything-model (SAM) [26]. Nonetheless, MF-HMI has a more complex data structure, and the boundary effects across AM frequencies are modality-specific, distinctive from noise patterns usually seen in other ultrasound images. Moreover, we aim to add complementary information to enhance segmentation using multi-modality images. For these reasons, we chose to train our network, HMINet [27], from scratch. The basic structure was adapted from *mmformer* [28], a multi-modality transformer-based network. Fig. 1c illustrates the architecture of HMINet. It has hybrid modality-specific encoders (**Conv encoder**: 5-stage convolutional encoder, **Transformer encoder**: 8 attention heads), a **modality-correlated encoder** (with Bernoulli indicators to randomly drop out a modality during training), and a 5-stage convolutional decoder (**Conv decoder**). The rationale for selecting *mmformer* as the backbone structure was based on several key characteristics: the hybrid, modality-specific encoding structure facilitated the local and long-distance context learning, and the following cross-modality encoder ensured feature alignments across B-mode and MF-HMI; the *auxiliary regularizer* encouraged additional independent modality learning, in order to prevent the model from biasing toward a specific modality and significant performance degradation when a modality was missing. We further incorporated a **channel-aware attention mechanism** [29] with both average-pooling and max-pooling in the **Conv decoder** to strengthen cross-channel learning, which is crucial for the multi-channel data in MF-HMI. To mitigate the domain differences between the training data and real clinical data, we applied a transfer learning strategy to fine-tune the model on clinical data and tested its performance using 4-fold cross-validation.

Several pre-processing steps were applied to input images: registration between MF-HMI images and the corresponding B-mode images; standardization (rescale to the range [0, 1]); for the training dataset, histogram equalization (*equalizeHist*) was applied to enhance the contrast; for phantom inclusions that were softer than the background and lesions monitored during FUS, where displacements within the inclusion were higher than the background, a linear transformation involving inversion and adding offsets, was applied to make all inclusion regions had lower values than the background. This approach reduced the false positive rate by preventing the network from misclassifying noisy regions as inclusions. All images were resized to 144 pixels x 144 pixels (0.07-0.13 mm per pixel). Data augmentation was applied during training, 50% of images were randomly cropped every epoch. For the FUS lesion, displacement images at later frames were subtracted from the first frame to show changes due to ablation. To accommodate different AM frequencies used for HMI and HMIgFUS, the order of MF-HMI images was not from low to high frequency but randomized every epoch, which also helped to restrain overfitting.

HMINet was trained for 250 epochs on NVIDIA RTX A5500, using an Adam optimizer (learning rate = 2e-4, betas = (0.9, 0.999), eps = 1e-8). The hybrid loss function is shown in (3), combining the area-based Dice Similarity Loss and distance-based Average Hausdorff Loss to increase sensitivity in margin detection and compensate for imbalanced size between the tumor and background pixels [30]. Segmentation accuracy was evaluated using the Dice Similarity Score (DSC), which, as shown in (4), measures the similarity of two sets of points. The network was first tested on phantom and mouse tumors where boundaries can be easily determined on B-mode through manual annotations. Later, we validated HMINet’s generalizability in *in vivo* human tumors and a FUS lesion.

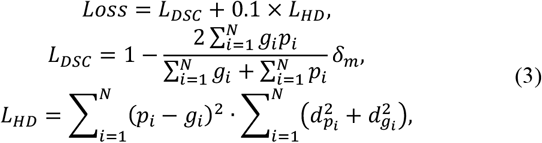

where *p*_*i*_ and *g*_*i*_ are the *i*th pixel in the network-segmented/reference boundary; 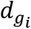 and 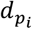 are the shortest distance between the *i*th pixel in the reference/network-segmented boundary to the network-segmented/reference boundary. The Dice Similarity Score is defined by:

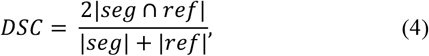

where seg and ref are the network-segmented/reference mask.

Reference boundaries of human breast tumors were delineated manually on research B-mode with the guidance of Butterfly B-mode and HMI images. Regions of interest (ROI) were generated automatically based on the HMINet-segmented boundaries for displacement ratio (DR) calculation. The tumor ROI covered 50% area of the bounding box of the segmented tumor, and the surrounding background ROI was at the same depth as the tumor ROI with a 2 mm lateral offset to the tumor boundary. DRs were calculated at all AM frequencies, and the median value was used to represent the stiffness at that time point. In FUS lesion segmentation, to provide output stability, boundaries were generated every four frames as the union of four masks derived from HMINet. Both B-mode and gross pathology were used as references to evaluate the network’s output. The overview pipeline is illustrated in Fig. 1d.

To assess the advantages of the multi-modality segmentation strategy, we compared the performance of HMINet using both modalities, B-mode-only and MF-HMI-only. By blinding the corresponding channels in the original architecture, we made comparisons on phantom, mouse, and clinical datasets.

## III. Results

### A. Segmentation of phantom inclusion

Fig. 3 (a) and (b) show representative B-mode, MF-HMI images from 100 to 1000 Hz, manually segmented and HMINet-segmented boundaries of a 2.5-mm, 70-kPa inclusion, and a 6.5-mm, 70-kPa inclusion. HMI images at 1000 and 700 Hz (the optimal frequencies with the highest contrast-to-noise ratio in each case) are overlaid on B-modes for visualization. The 2.5-mm, 70-kPa inclusion has better detectability in high AM frequencies since a small inclusion in HMI was more prone to axially shearing effect; the 6.5-mm, 70-kPa inclusion can be clearly seen across all AM frequencies. Reduced boundary effects are observed as AM frequencies increase due to less shearing in the axial direction. Segmentation results using different inputs are summarized in Table 1. Each DSC score was averaged over 8 independent measurements acquired at different sequences and stiffness.

**TABLE I.** Patient summary.

| Age | Molecular subtype | Stage |
| --- | --- | --- |
| Patients with NACT |  |  |
| 57 | ER-/PR-/HER2- | IIB |
| 53 | ER+/PR+/HER2- | IIIB |
| 81 | ER+/PR-/HER2- | IIB |
| 62 | ER-/PR-/HER2- | IIB |
| 38 | ER+/PR-/HER2+ | IIB |
| Patients without NACT |  |  |
| 74 | ER+/PR+/HER2- | IA |
| 71 | ER+/PR+/HER2- | / |
| 67 | ER+/PR-/HER2- | / |

**TABLE II.** Segmentation results for phantom studies (N = 8)

| Diameter (mm) | Multi-modality | B-mode only | HMI only |
| --- | --- | --- | --- |
| 2.5 | **0.87** | 0.78 | 0.85 |
| 4.1 | **0.91** | **0.90** | 0.76 |
| 6.5 | **0.95** | **0.94** | 0.84 |
| 10.5 | **0.96** | **0.94** | 0.90 |
| Average | **0.92** | 0.89 | 0.84 |

**Fig. 3.**
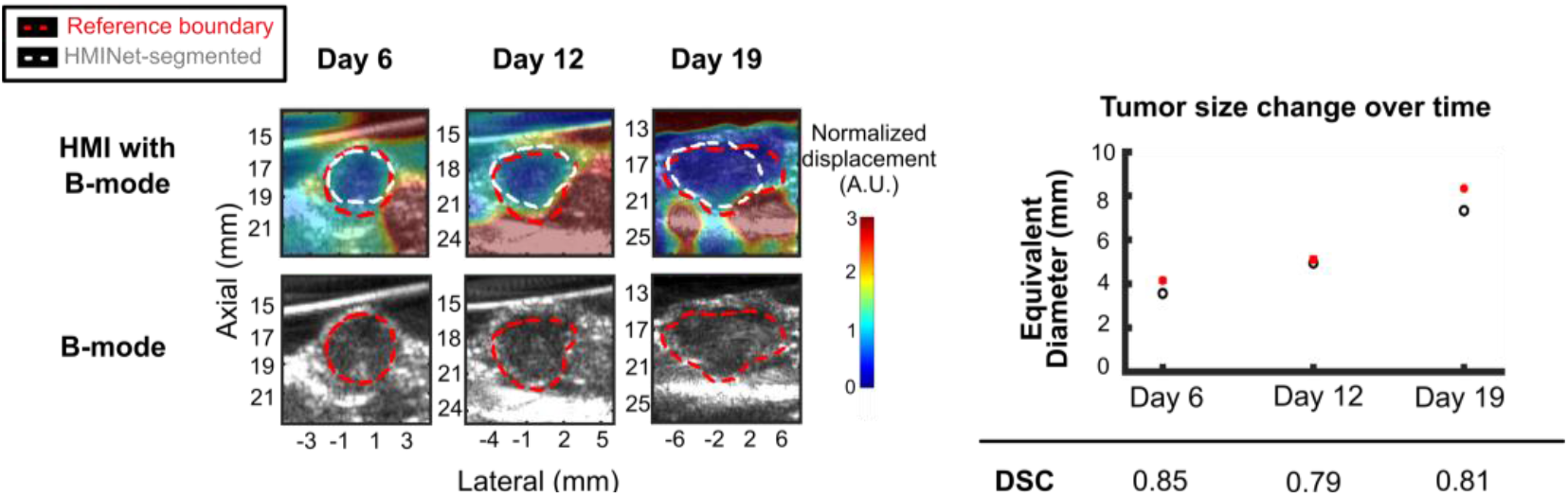
*In vivo* 4T1 murine tumors with HMINet-segmented (white) and manually segmented boundaries (red).

The multi-modality (MF-HMI and B-mode) network performed the best in all cases, with a significantly higher average DSC value of 0.92, compared with a DSC value of 0.89 in the HMI-only and 0.84 in the B-mode-only network. Segmentation accuracies increase with the inclusion size, and the best performance is achieved with the 10.5-mm inclusions, as both B-mode and MF-HMI obtained better image quality for larger inclusions. Additionally, B-mode images of the 2.5-mm inclusions had insufficient signal-to-noise ratio (SNR), leading to ambiguity in the reference boundary determination and subsequently affecting the performance evaluation.

### B. Monitoring of murine tumor progression

A representative case of longitudinal monitoring of murine tumor progression using MF-HMI is shown in Fig. 3. HMI images with the highest CNR are overlaid on B-mode images for visualization. It is observed that the tumor grew larger and stiffer over time. HMINet performed well in all time points with an average DSC of 0.85, 0.79, and 0.81. Tumor size change was calculated, and the equivalent diameter computed based on area was plotted against time.

### C. Assessment of human breast tumor response to NACT

Fig. 4 shows the segmentation and NACT prediction results in four clinical cases. The pretreatment DRs of patients 1 to 4 were 0.43, 0.11, 0.13, and 0.20, indicating that patients 1 and 4 had slightly softer tumors compared to patients 2 and 3. Three weeks later, the DR of patients 1 and 3 decreased, which suggested tumor stiffening and not responding. Conversely, significant increases (345% and 160%) in DRs were observed in patients 2 and 4. Based on these DR changes, patients 1 and 3 were predicted as npCR, which were later validated by the pathology results after surgery. Patients 2 and 4 were predicted to achieve pCR (patient 2 is still pending pathological confirmation). It is worth noting that at only three weeks into NACT, when the tumor size hadn’t changed much, if any, HMI-derived DR was sensitive enough to predict the pathological result at the treatment endpoint. Interestingly, as shown in the 3^rd^ row in Fig 4, segmentation using only B-mode images led to certain errors. Poor performance was prominent in patients 2, 3, and 4 when the tumor had fuzzy boundaries and mixed echogenic patterns, potentially due to multi-layers and calcification (appearing as bright clusters in B-mode). Similar findings were also seen in other automated B-mode-based breast tumor segmentation methods [31]. Calcification, i.e., the deposition of calcium, produces large echoes that could be problematic in B-mode-based segmentation, but it doesn’t substantially impact elasticity imaging due to the small portion commonly seen in malignant tumors [32]. The misleading fine structures were segmented correctly by adding MF-HMI, as shown in the 2^nd^ row.

**Fig. 4.**
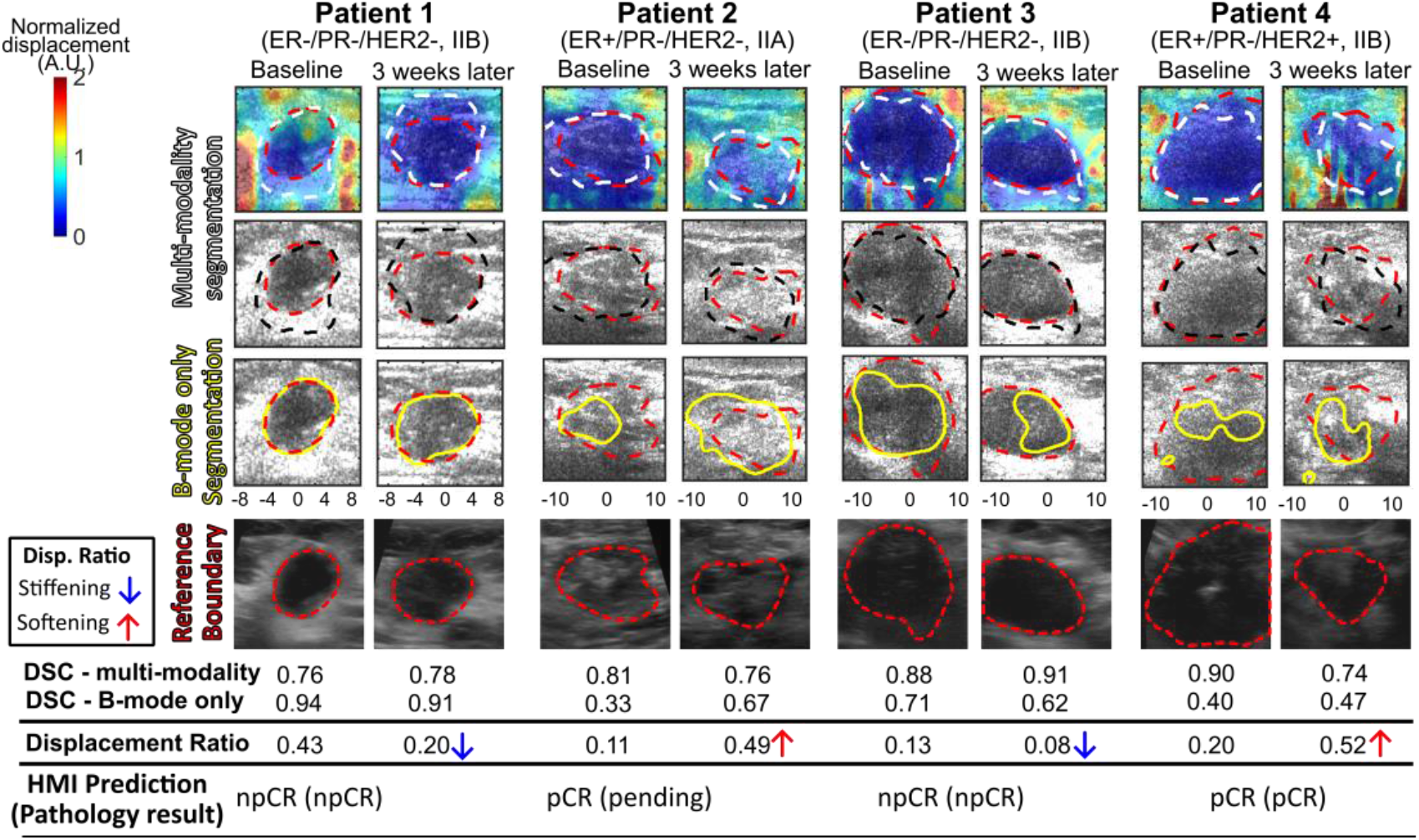
Breast tumor segmentation and the following automated NACT prediction. Reference boundaries (red), HMINet-segmented boundaries using multi-modality (white/black, shown in 1^st^ and 2^nd^ row) and B-mode only (yellow, shown in 3^rd^ row). NACT predictions were made by comparing displacement ratios at three weeks with those at baseline.

### D. Monitoring of FUS lesioning

Fig. 5 shows the HMINet-segmented FUS-induced lesion on MF-HMI and B-mode images during ablation. Though the FUS lesion gets stiffer due to protein denaturation after ablation, tissue displacements around the FUS focus were higher than the unabated region, potentially due to the liquifying process under heat deposition. It is observed that at 550 Hz, the perceived lesion size was smaller compared to 150 Hz; however, the contrast was not as high as that at 150 Hz. Though B-mode before and after FUS ablation did show some differences, the lesioning process could not be observed during ablation. As shown in Fig. 5, MF-HMI was able to monitor the lesioning progress. Gross pathology was performed on the imaging plane immediately after ablation; a reference boundary was drawn and registered to HMI/B-mode images, and the DSC was calculated to be 0.87. The lesion boundary was also delineated from the difference between B-mode images before and after ablation, as shown within the blue line.

**Fig. 5.**
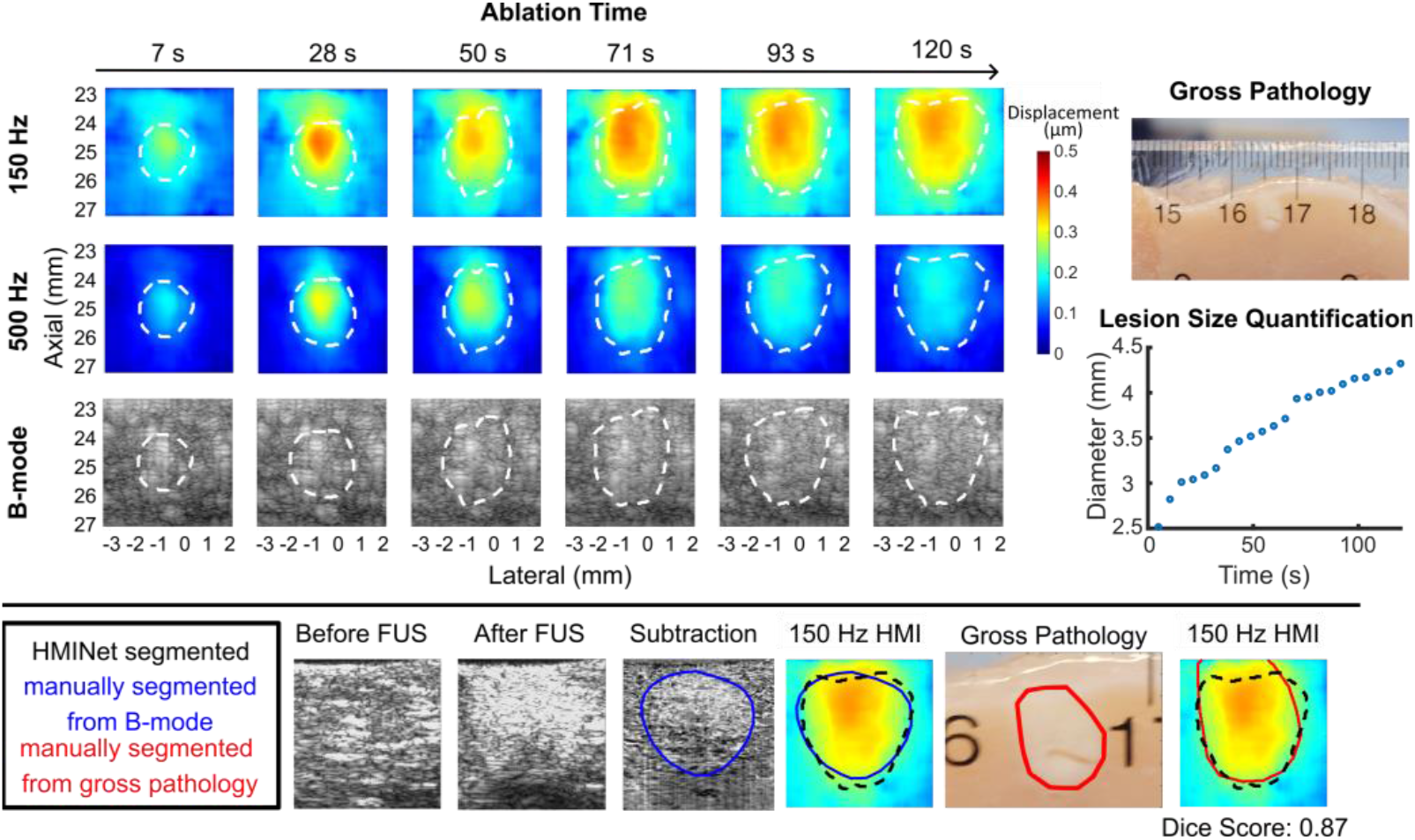
FUS-lesion monitored by HMIgFUS system on *ex vivo* chicken tissue. HMI images at 150 Hz and 500 Hz and B-mode images during ablation were shown.

### E. Network comparison

To assess the effectiveness of multi-modality and transfer learning strategy in enhancing clinical applicability, we compared the performance of networks under different conditions: trained by both MF-HMI and B-mode images with or without transfer learning, trained by HMI only, trained by B-mode only. In Fig.6, Kruskal-Wallis tests revealed statistical differences in network performance across phantom (n = 32), mouse (n = 24) datasets, and clinical datasets (n = 16). Two observations are noteworthy: in mouse and phantom datasets, the B-mode-only network performed better than the HMI-only network, as the echogenicity patterns were quite simplified, and inclusions had well-defined boundaries and clean backgrounds. Under these circumstances, segmentation inaccuracies mainly came from boundary effects. In clinical datasets, however, where tumor boundaries were ambiguous and the physiological properties of tissue were influential, the B-mode-only network failed to produce quality results consistently. The HMI-only network performed significantly better than B-mode-only, and the best performance was achieved by the multi-modality network with transfer learning, with an average DSC (over ten repeated predictions) of 0.821, followed by HMI with transfer learning with a DSC of 0.819, and HMI without transfer learning with a DSC of 0.800. The comparison empirically demonstrated that combining both imaging modalities and transfer learning improved the network’s segmentation robustness in clinical scenarios.

**Fig. 6.**
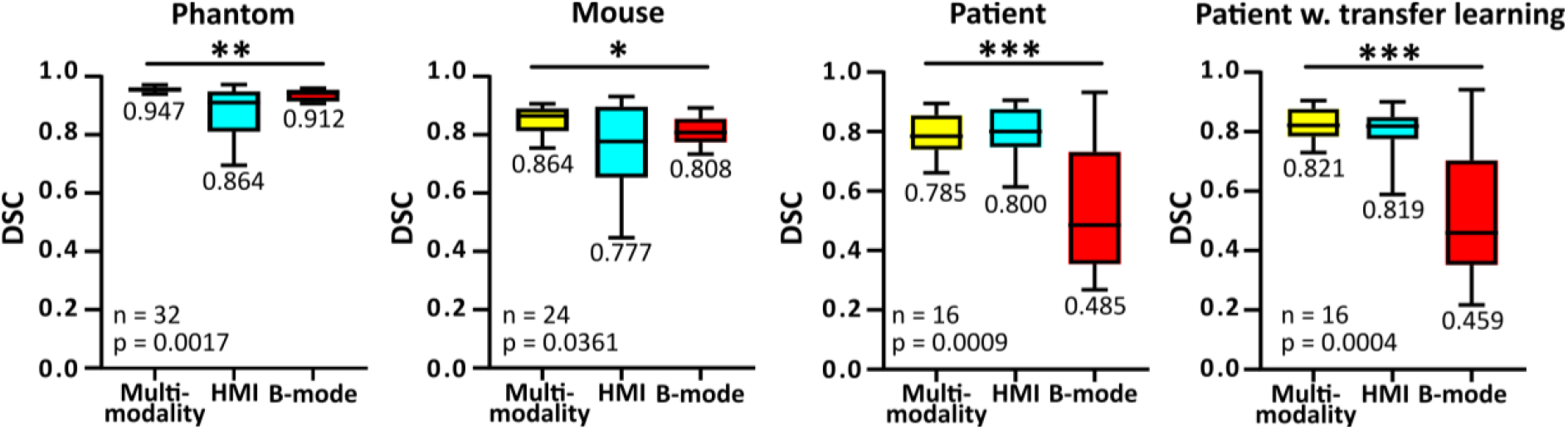
Kruskal-Wallis tests of networks’ performance on phantom, mouse, and patient datasets. Significant differences in segmentation accuracy were shown across networks using different imaging modalities (MF-HMI only, B-mode only, and both), with/without transfer learning.

## IV. Discussion

HMI is an elasticity imaging technique by inducing localized, oscillatory tissue movement, and the resulting on-axis displacement is estimated during excitation. It is operator-independent and provides deep penetration up to a focal depth of 30 mm [17]. Stiffness alterations in tumors can occur prior to any noticeable changes in anatomical changes, which grants HMI the potential to make early predictions of NACT treatment response. At the same time, because of its experimental design, it offers real-time FUS lesioning monitoring with excellent image quality that allows for precise margin delineation to guarantee complete therapy.

In this study, we developed an automated tumor and FUS lesion assessment method for multi-frequency HMI systems. The goal was fully utilizing MF-HMI (different boundary effects) and the multi-modality information (acoustic and mechanical) from B-mode and MF-HMI. We explored this idea using neural networks and further improved its clinical performance using transfer learning. HMINet was shown to achieve high segmentation accuracy and predicted tumor response to NACT based on stiffness changes as early as three weeks into treatment. Furthermore, networks with different inputs were compared, which empirically validated the benefits of combining imaging modalities. It is also worth emphasizing that the method we proposed has actual real-time potential for FUS therapy monitoring. After training HMINet in PyTorch, HMINet can be exported as an Open Neural Network Exchange (ONNX) format model, which can be integrated with the HMIgFUS system in MATLAB for real-time application. Based on our preliminary test, the pre-trained, lightweight version of HMINet produced a segmentation frame rate of 0.2 Hz.

In MF-HMI, DR is influenced not only by frequency-dependent boundary effects due to the shearing in the axial direction after external acoustic radiation force excitation but also by the internal viscoelasticity of targeted tissue. As a result, median values of DR were used to assess stiffness change over time to avoid a biased analysis due to the less satisfactory quality of a specific AM frequency. In future work, the DR-AM frequency relationship will be investigated to correlate with the viscoelastic properties of tumors, which can potentially serve as another biomarker for tumor response prediction besides DR.

In the proposed network, the input order of frequency was randomized. Though the effects of AM frequency on HMI image quality have been explored in phantoms, the relationship hasn’t been fully established in mouse and patient data. By shuffling the frequency order, we aimed to force the network to make decisions based on the most significant features in those multi-channel images but not the trends across channels. It reduces the risks of network overfitting that is biased toward phantom patterns or particular HMI sequences and promotes the transferability of the network in different AM sequences.

Limited clinical data has always drawn concerns in the field of medical imaging about the network’s domain adaptation and failure in unseen cases. In this study, we imaged phantoms covered by *ex vivo* tissues and wire mesh to introduce phase aberrations and mimicked clinical data acquisition. The segmenting ability of HMINet in murine and human tumors demonstrated that, despite the network being trained on circular phantoms, it wasn’t biased by circular shapes and remained flexible in predicting irregular, less circular shapes.

Determining reference boundaries in clinical and FUS lesion datasets could be challenging as there was no actual ground truth. In the future, histopathological analysis will be performed to compare the tumor’s largest dimension of the tumor with that derived from the predicted boundaries. Intriguingly, in the first clinical case (in Fig. 4a), the predicted tumor size was larger than its perceived size in B-mode because of larger tumor sizes observed in HMI images. Similar results were observed by Farrokh *et al*., they showed that B-mode has a higher risk of size underestimation by comparing the size measured by B-mode, shear wave elastography with histopathological tumor size in invasive lobular carcinoma (ILC) [33], Hall *et al*. demonstrated larger areas of lesion size in strain images than B-mode in invasive ductal carcinoma (IDC) [34]. In contrast to benign lesions, malignant tumors are associated with desmoplasia reactions where the excess growth of fibroblasts causes stiffening in the perilesional area and can be captured by elasticity imaging [35]. A limitation of our work is that the training dataset didn’t include cases where inclusions have larger sizes in elasticity than B-mode imaging. In the next step, we will increase our training dataset by adding more cases of *ex vivo* malignant tumors. We will also validate our findings by obtaining HMI images of *in vivo* benign tumors, where less prominent perilesional stiffening is expected.

In addition to improving segmentation accuracy, gaining a deeper understanding of the impact of AM frequency on boundary detectability holds great clinical value. Image quality assessment criteria, such as generalized contrast-to-noise ratio (gCNR), which is proportional to the maximum separation success probability and thus robust against dynamic range alteration, will be considered for ranking the significance of AM frequencies [36]. Following that, a selected set of HMI images will be used for boundary segmentation instead of the full range of AM frequencies. Besides, deep learning-based interpretation methods will be used, such as the GradCam-generated heat maps [37] and activation maps [38], to identify the most spatially relevant regions in the image and the associated dominant AM frequencies. We could further validate their significance by observing the performance change in the absence of one or several AM frequencies.

From the perspective of data acquisition, the image quality of MF-HMI could be further improved. Liu et al. demonstrated that the displacement tracking sequence can be optimized in conventional HMI [39]. To improve image quality, especially in clinical data, imaging parameters such as the pressure of ARF, parallel tracking, and F-number can be further investigated.

Moving toward the goal of predicting NACT efficacy and facilitating real-time FUS ablation monitoring, this automated segmentation network will serve as the initial stage for tumor and FUS lesion quantification. In the future, we will increase clinical data and establish other HMI-derived biomarkers, such as perilesional stiffening area, for more precise, subtype-specific NACT prediction. We will also systematically verify its robustness on FUS lesion segmentation by testing on more FUS lesions.

## V. Conclusion

In this study, we developed an automated tumor and FUS lesion quantification method using a multimodality, transformer-based network, HMINet. It exhibited superior performance on phantom, *in vivo* mouse, patient, and FUS-lesion datasets compared with single modality networks, achieving an average DSC score of 0.95, 0.86, 0.82, and 0.87, respectively. Displacement ratios were calculated based on HMINet-segmented boundaries to predict tumor responses to NACT. Future work will be focused on deepening our understanding of the impact of AM frequency on lesion detectability and developing a physiology-informed model for NACT prediction and individualized treatment planning.

## Acknowledgments

This study was supported by the National Institutes of Health (R01CA228275). The authors thank Julia E. McGuinness, M.D., at Columbia University Irving Medical Center, Murad Hossain, Ph.D., Tuhin Roy, Ph.D., Melina Tourni, M.S., Chunqi Li, Ph.D., for their constructive discussion, and Pablo Abreu, M.A., for his administrative assistance.

## Notes

This work was supported in part by the National Institutes of Health (R01CA228275).

### Competing Interest Statement

The authors have declared no competing interest.

